# VSS-Hi-C: Variance-stabilized signals for chromatin contacts

**DOI:** 10.1101/2021.10.19.465027

**Authors:** Neda Shokraneh Kenari, Faezeh Bayat, Maxwell Libbrecht

## Abstract

**Motivation:** The genome-wide chromosome conformation capture assay Hi-C is widely used to study chromatin 3D structures and their functional implications. Read counts from Hi-C indicate the strength of chromatin contact between each pair of genomic loci. These read counts are heteroskedastic: that is, a difference between the interaction frequency of 0 and 100 is much more significant than a difference between the interaction frequency of 1000 and 1100. This property impedes visualization and downstream analysis because it violates the Gaussian variable assumption of many computational tools. Thus heuristic transformations aimed at stabilizing the variance of signals like the shifted-log transformation are typically applied to data before its visualization and inputting to models with Gaussian assumption. However, such heuristic transformations cannot fully stabilize the variance because of their restrictive assumptions about the mean-variance relationship in the data.

**Results:** Here we present VSS-Hi-C, a data-driven variance stabilization method for Hi-C data. We show that VSS-Hi-C signals have a unit variance improving visualization of Hi-C, for example in heatmap contact maps. VSS-Hi-C signals also improve the performance of subcompartment callers relying on Gaussian observations. VSS-Hi-C is implemented as an R package and can be used for variance stabilization of different genomic and epigenomic data types with two replicates available.

**Availability:** https://github.com/nedashokraneh/vssHiC

## 1 Introduction

The 3D organization of chromatin in the nucleus plays a key role in many cellular processes. Chromatin conformation can control transcription through mechanisms including positioning genes within nuclear components—such as nuclear speckles—enriched with transcription factors Chen *et al*. (2018), organizing genomic elements into topological association domains (TADs) Stadhouders *et al*. (2019), colocalizing genes and their enhancers, and facilitating repression Szabo *et al*. (2019). The genome-wide chromosome conformation capture assay Hi-C assesses chromatin conformation by measuring the frequency of contacts or interactions between pairs of genomic bins. Inferring biological insights from Hi-C data requires accurate computational analysis methods.

Researchers have developed computational tools for processing Hi-C data into a format useful for downstream analysis, including tools for read aggregation (Durand *et al*., 2016; Abdennur and Mirny, 2020) and bias normalization (Imakaev *et al*., 2012; Knight and Ruiz, 2013). These tools output a matrix of contact counts that measures the contact strength of each pair of genomic positions. However, similar to other sequencing-based measurements, these signals are heteroskedastic; that is, a difference between the interaction frequency of 0 and 100 is much more significant than a difference between the interaction frequency of 1000 and 1100 (Motakis *et al*., 2006; Anders and Huber, 2010; Ahlmann-Eltze and Huber, 2023) (defined precisely below).

This property of heteroskedasticity impedes visualization and downstream analyses. For example, a heatmap visualization of raw Hi-C counts has its color map dominated by large outliers, making it impossible to see subtle contact patterns important for loops and compartments. Likewise, many existing analysis methods (Yaffe and Tanay, 2011; Rao *et al*., 2014) assume signals follow a Gaussian distribution because doing so is easy to implement, but this assumption is inaccurate for heteroskedastic signals. The same issue also afflicts existing machine learning models that minimize mean-squared error (Zhang *et al*., 2018; Dimmick, 2020), as doing so is mathematically equivalent to maximizing the log-likelihood of a Gaussian model. Also, some statistical methods for calling TADs rely on Gaussian data (Lévy-Leduc *et al*., 2014) and variance-stabilizing transformation is necessary before applying these methods.

Two approaches have been proposed to handle heteroskedastic signals. One strategy to address heteroskedasticity is to use a non-Gaussian model, such as a negative binomial distribution, which explicitly accounts for the mean-variance relationship. However, such models can be hard to implement and optimize. Instead, many existing approaches first aim to stabilize variance with a transformation, such as a heuristic shifted log (log(*x*+1)) and asinh 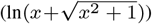. However, such transformations assume a specific mean-variance relationship (e.g. log assumes a quadratic relationship) and thus do not always successfully stabilize variance (Ahlmann-Eltze and Huber, 2023). (See section 2.4 for a review of all previously proposed transformations.)

Thus transformation methods for other data types typically apply a data-driven variance-stabilizing transformation. For example, researchers typically stabilize the variance of RNA-seq using the data-driven variance-stabilizing transformation approach in DESeq (Anders and Huber, 2010). Similarly, we recently proposed the delta method-based approach VSS (Bayat and Libbrecht, 2021) that stabilizes the variance of 1D genomic signals such as ChIP-seq.

Here, we propose a method, VSS-Hi-C, for stabilizing the variance of Hi-C contact signals. Our contributions presented here are as follows. First, we adapted the VSS method to extend it to handle 2D Hi-C contact matrices. Second, we present a user-friendly R package that provides an interface with an easy-to-use class (InteractionSet) and data format (.cool) for Hi-C data. Third, we propose three evaluation metrics that measure the downstream impact of heteroskedasticity on Hi-C analysis and benchmarked all approaches for Hi-C signals. We found that VSS-Hi-C signals have more stable variance compared to all other transformations and produce informative visualization of chromatin structures at different scales, and we found that VSS-Hi-C signals outperform or perform comparably to other transformations in terms of identifying more biologically validated subcompartments.

## 2 Materials and Methods

Our data sources are provided in Table. S1.

### 2.1 Problem definition

The read counts from the Hi-C assay represent interaction frequencies between pairs of genomic bins. As mentioned above, these read counts do not have constant variance, meaning we expect an observed interaction between a pair of genomic bins with an observed interaction of 1000 reads to have different (usually higher) uncertainty than another pair of genomic bins with 100 reads. VSS-Hi-C utilizes a Hi-C experiment with two biological replicates to estimate the mean-variance relationship in the experiment (section 2.3) and derive a variance-stabilizing transformation (section 2.2).

More formally, the inputs are two Hi-C contact matrices *M* ^1^, *M* ^2^ *∈* ℕ^*n×m*^ corresponding to interaction frequencies between chromosomes *c*1 and *c*2 from two biological replicates (*n* and *m* are the number of genomic bins in chromosomes *c*1 and *c*2 respectively). VSS-Hi-C estimates the mean-variance relationship in data (Fig. 1b), derives a variance-stabilizing transformation *t*(*x*) (Fig. 1c) and outputs transformed contact matrices *t*(*M* ^1^) and *t*(*M* ^2^) which have a unit variance; that is var(*M*_*i,j*_) = 1 regardless of the magnitude of *M*_*i,j*_ (Fig. 1a-2a).

**Fig. 1:**
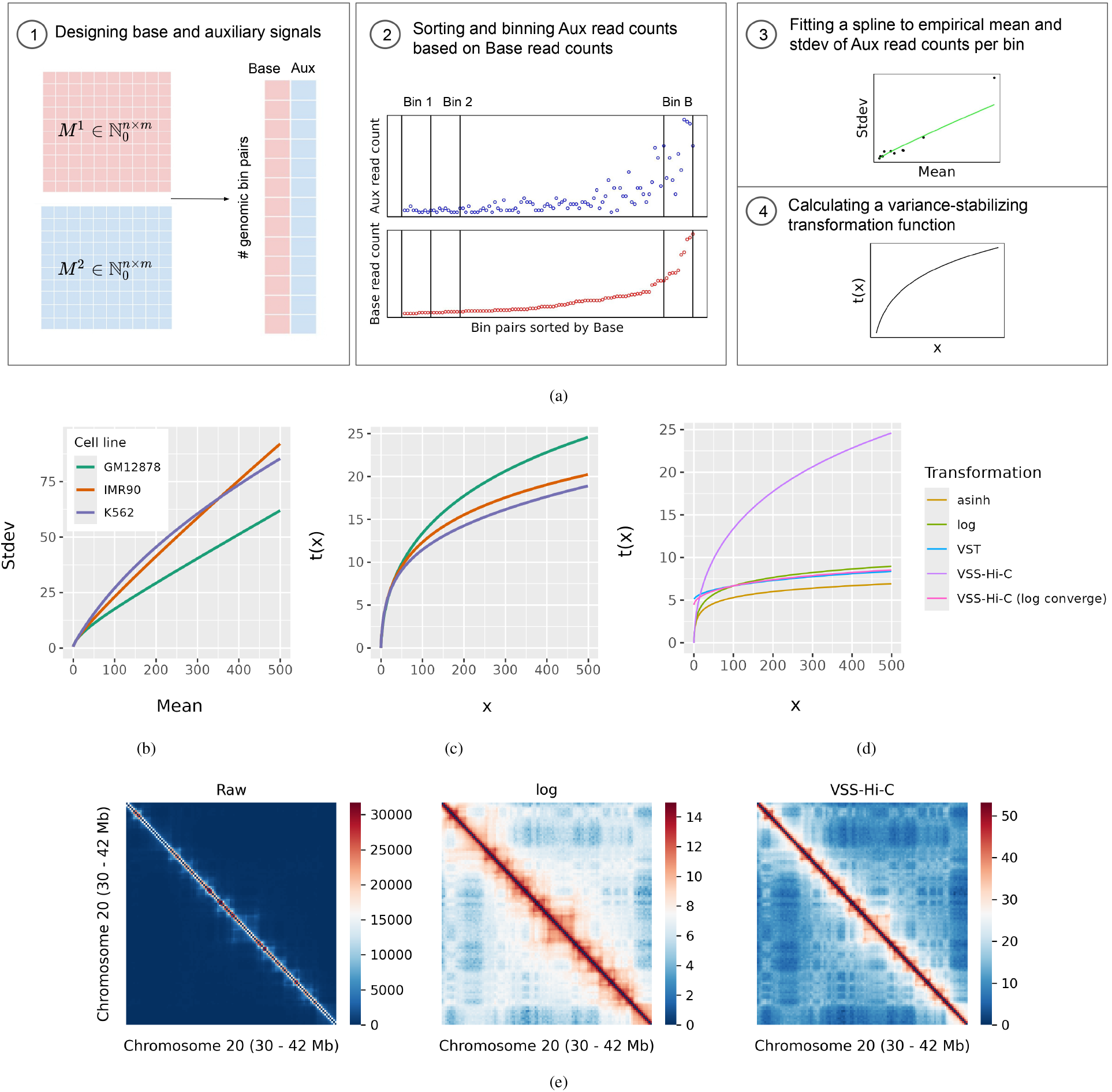
(a) Overview of VSS-Hi-C for variance stabilization of Hi-C data. (b) Learned mean-stdev curves for three Hi-C experiments, each from a different cell line. (c) Corresponding VSS-Hi-C transformations to mean-stdev curves in (b). (d) Comparison of different transformation methods. (e) Comparison of heatmap visualization of raw, log-transformed and VSS-Hi-C signals with linear colour scale. (d) and (e) represent transformation functions and heatmap visualizations of the GM12878 cell line.

**Fig. 2:**
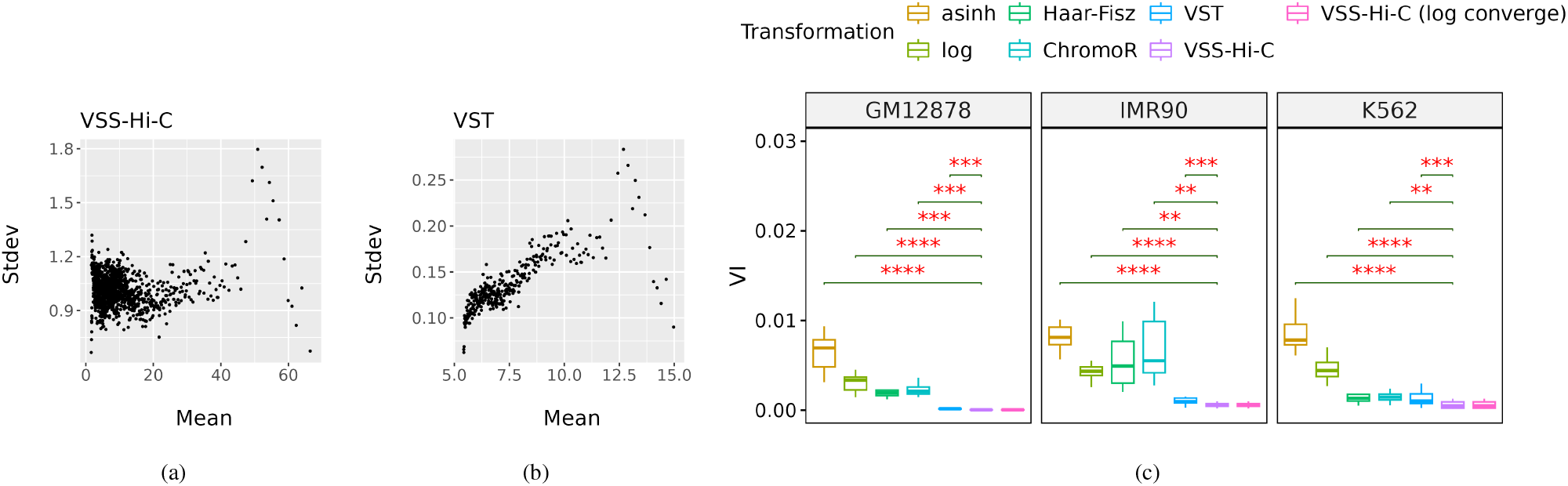
(a,b) Standard deviation (stdev) vs mean for VSS-Hi-C and VST signals respectively. These plots are for chromosome 13 of the GM12878 cell line. (c) Variance instability (VI) of transformed signals for 3 cell lines. Each box represents VIs for chromosomes 13 to 22. Red asterisks indicate the significance of paired one-sided t-tests with an alternative hypothesis: VSS-Hi-C signals have less VI compared to other transformed signals. Non-significant labels (*p >* 0.05) are removed. 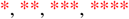 represent *p* less than 0.05, 0.01, 0.001, and 0.0001 respectively.

VSS-Hi-C stems from a general idea for variance-stabilizing transformation, a delta method. First, we briefly explain a delta method and then describe VSS-Hi-C in detail.

### 2.2 Variance-stabilizing transformation based on the delta method

The delta method is a general method for approximating a probability distribution of a function of an asymptotically normal random variable with a known variance. Let *X* be a random variable with E[*X*] = *μ* and Var[*X*] = *σ*^2^. A delta method approximates the mean and variance of *Y* = *g*(*X*) where *g* is any function that its first derivative at *μ, g*^*t*^(*μ*), exists and is non-zero. A first-order Taylor approximation for *Y* = *g*(*X*) is:

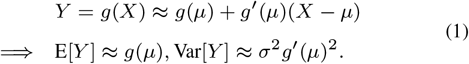

Considering there is a known relationship between the mean and variance of a random variable (E[*X*] = *μ* and Var[*X*] = *h*(*μ*)), the goal is to find a function *g* such that *Y* = *g*(*X*) has a variance independent of *μ*. Forcing the condition Var[*Y*] ≈ *h*(*μ*)*g*^*t*^(*μ*)^2^ = *C*, where *C* is a constant, implies a differential equation:

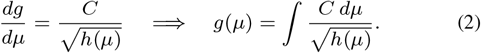

Thus we can construct a variance-stabilizing transformation function given any known mean-variance relationship. There are well-known approximate variance-stabilizing transformations for probability distributions with known mean-variance relationships. For example, the negative binomial (gamma-Poisson) distribution commonly used to model RNA-seq and scRNA-seq counts has a quadratic mean-variance relationship Var[*Y*] = *μ* + *αμ*^2^, where *μ* and *α* are the mean and overdispersion of the distribution respectively. Applying a delta method implies an approximate variance-stabilizing transformation *g*(*y*) = log(*y* + *y*_0_), which is the widely-used shifted logarithm transform.

To avoid assuming a specific mean-variance relationship, here we estimate the mean-variance relationship empirically and use equation 2 to find a variance-stabilizing transformation *g*, following the previously-described method VSS for 1D genomic data sets (Bayat and Libbrecht, 2021) (section 2.3).

### 2.3 VSS-Hi-C

VSS-Hi-C has 4 steps to find a data-driven variance-stabilizing transformation (Fig. 1a):

Step 1: We designate *M*_1_ as the “base” replicate and and *M*_2_ as “auxiliary” replicate and design a two-column matrix as follows. For intra-chromosomal matrices (*c*1 = *c*2), we flatten all the elements in and above the main diagonal of matrices and for inter-chromosomal contact matrices *c*1 =*/ c*2, we flatten whole matrices to fill the base-aux matrix. Each row of the base-aux matrix represents two observations of one pair of genomic bins (Fig. 1a-1).

Step 2: The purpose of constructing a base-aux matrix is to find pairs of genomic bins whose interaction frequencies are likely from the same distribution. For that purpose, we use base observations to order the base-aux matrix, group aux observations into equal-size bins, and assume that pairs of genomic bins within each bin have a similar variance for the purposes of learning (Fig. 1a-2).

More formally, define 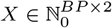 as a base-aux matrix where *BP* is a number of genomic bin pairs. We sort *X* based on its first column (base observations) into 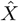. Given a bin size of *s*, we group the rows of 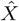 into 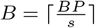 bins.

In VSS-Hi-C, we only use 2 replicates and choose one replicate as a base and another as an auxiliary signal. This can be changed to support more than 2 replicates similar to VSS, however, we have not found any Hi-C experiment with more than 2 replicates.

Step 3: Assuming that aux read counts within one bin are samples from one distribution, we calculate an empirical mean and variance of such distribution for bin *b* as:

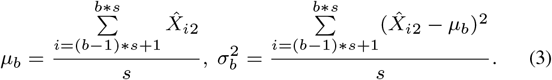

Note that in the rest of the text and all figures, we show standard deviation or stdev (*σ*) instead of variance (*σ*^2^) as it is an essential statistic for the estimation of a variance-stabilizing transformation (section 2.2).

Given *B* mean and stdev estimations for *B* bins, we fit a spline to 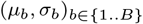 pairs to estimate the mean-stdev relationship, 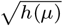.

Step 4: Given the estimated mean-stdev relationship *h* from step 3, we estimate the variance-stabilizing transformation *t* based on equation 2. Since *h* is not parametric, *t* is obtained by the numerical integral of *h*(*x*) over a set of *X* along a range of signals (asinh-spaced).

### 2.4 Alternative transformations

Here we briefly describe alternative strategies for transforming Hi-C signals for the purpose of variance stabilization.

#### 2.4.1 log and asinh

When the mean-variance relationship is *h*(*μ*) ∝ *μ*^2^, the variance-stabilizing transformation is proportional to the log function (Ahlmann-Eltze and Huber, 2023). This transformation is widely used to preprocess bulk and single-cell genomic data. We use a log transformation (log(*x*+1)) as a heuristic alternative. Another heuristic transformation is a hyperbolic arcsine function asinh 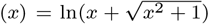 which is defined for all real values and is a popular substitute for shifted log transform.

#### 2.4.2 ChromoR (Haar-Fisz transform)

ChromoR (Shavit *et al*., 2014) is a computational tool for Hi-C data that employs Haar-Fisz transformation (Supplementary section 1.1) and wavelet shrinkage methods to stabilize the variance and reduce the noise of observations, respectively. We use ChromoR (Haar-Fisz + denoising) and Haar-Fisz in our evaluations. The Haar-Fisz transform uses data from a single replicate, employing the assumption that neighboring genomic positions are pseudo-replicates of one another.

#### 2.4.3 VST

An alternative data-driven variance-stabilizing transformation is a vst function from DESeq2 package (Love *et al*., 2014) originally designed for RNA-seq gene expression data. This tool estimates per-locus-pair mean and dispersion (Supplementary section 1.2) and estimates the mean-variance relationship; like VSS-Hi-C, it uses this curve to derive a variance-stabilizing transformation based on the delta method. We found that VST does not fully stabilize variance, perhaps due to producing separate estimates for each locus pair, requiring more replicates that are available for Hi-C data (Results).

#### 2.4.4 VSS-Hi-C (log converge)

In order to place transformed signals in a familiar scale, the vst function from DESeq2 linearly scales the estimated transformation function to converge to the log function asymptotically. More formally, given a variance-stabilizing transformation function *t*(*x*), they define *t*^*′*^(*x*) = *a × t*(*x*) + *b* such that *t*^*′*^(*x*) = *log*(*x*) for large *x* values. Then, they use two *x* values, *x*_1_ = quantile(*X*, 0.95) and *x*_2_ = quantile(*X*, 0.99999) (X is a set of raw signal values), to define a system of equations *t*^*′*^(*x*_*i*_) = *a × t*(*x*_*i*_) + *b* = *log*(*x*_*i*_) and find *a* and *b*. We employ a similar strategy and use VSS-Hi-C converging to the log function named VSS-Hi-C (log converge) in our evaluations. While the resulting units have a familiar scale and retain variance-stability, they lose the property of unit variance (Fig. S1e) and we found they perform slightly worse when input to downstream tools (Results).

### 2.5 Evaluation metrics

#### 2.5.1 Variance instability

To evaluate whether a given transformation method successfully stabilizes variance across different signal strengths, we employ the same binning strategy in VSS-Hi-C to group signals. We define the variance instability (VI) metric as the variance of variance of signals in each group. Therefore, more instability of variance results in higher VI.

More formally, given transformed signals, 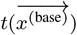 and 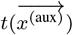, we use the same binning strategy introduced in section 2.3 to group 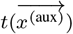 signals into *B* bins. Note that ordering by 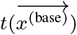 is the same as ordering by 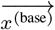 since *t* is a monotonic function. Assuming *I*_*j*_ be the set of positions in bin *j*, we define 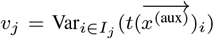 as a variance of signals within bin *j*. Finally, we define the VI metric as the scaled variance of *v*_*j*_ across bins,

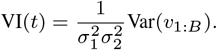

The 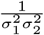 is a normalization factor to enforce *t*(*x*) and *αt*(*x*) have the same VI value for a constant *α. V I*(*t*) equals zero when the empirical variance is equal across all signal magnitudes; higher *V I*(*t*) indicates worse residual variance instability.

#### 2.5.2 Downstream analyses

##### Compartment analysis

The chromatin is segregated into two or more states ((sub)compartments), each enriched with specific properties like epigenomic, replication timing, and transcription patterns. Originally, two compartments, A and B, were identified based on their preferential interaction with each other (genomic bins within A compartment interact mostly with other genomic bins in A, and vice versa) (Lieberman-Aiden *et al*., 2009). (Rao *et al*., 2014) shows that two compartments can be further divided into different subcompartments with different epigenomic patterns by analysis of higher-resolution Hi-C data. They cluster genomic bins into subcompartments according to their long-range (inter-chromosomal) interaction profiles by the Hidden Markov model (HMM).

Following (Rao *et al*., 2014), we use chromosomes 1 to 4 to create odd-even inter-chromosomal contact matrix, 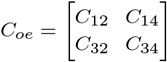, where *C*_*ij*_ is inter-chromosomal contact matrix between chromosomes *i* and *j*. Note that rows and columns of *C*_*oe*_ include genomic bins within odd and even chromosomes, respectively. Then, we apply a hidden Markov model (HMM), a clustering method with stable solution with Gaussian distributed variables, on *C*_*oe*_ and its transpose 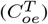 to annotate odd and even chromosomes respectively. (For comparison, we also tried a KMeans clustering method.) Finally, we biologically evaluate subcompartment annotations to assess the impact of transformations on finding more biologically meaningful subcompartments. To biologically evaluate the subcompartment annotation, we hypothesize that positions in the same subcompartment should have similar transcription factor binding and histone modification activity. Thus, following previous work (Libbrecht *et al*., 2015; Shokraneh *et al*., 2023), we calculate the variance explained metric (VE), a fraction of the variance of a relevant signal that is explainable by annotation. This metric is bounded by [0, 1], and higher values indicate more agreement between annotation and signal values. We use 12 epigenomic signals including ChIP-seq targeting H2A.Z, 10 histone modifications and DNase-seq.

##### TAD analysis

Topologically associating domains (TADs) are diagonal blocks in Hi-C contact matrices representing self-interacting genomic regions. They have important functional roles like they are linked to gene regulation by exposing and insulating regulatory elements to gene promoters (Symmons *et al*., 2016). Different algorithms exist to identify TADs from Hi-C contact matrices taking raw or normalized data as input (Zufferey *et al*., 2018). Overall, TAD callers are based on linear scores, statistical models, network features or clustering. We use one TAD caller per category to assess the performance of these algorithms given transformed data: (1) TopDom (Shin *et al*., 2016) is based on linear scores and had the best overall performance in a benchmarking study (Zufferey *et al*., 2018), (2) SpectralTAD uses spectral clustering, it is efficient and its detected domain boundaries are enriched with biological features (Cresswell *et al*., 2020), and (3) HiCseg (Lévy-Leduc *et al*., 2014) is a statistical model which was also among the top scoring in a benchmarking study (Zufferey *et al*., 2018).

We apply TAD callers on raw and transformed data to assess the validity of identified TADs after different transformations. Following a previous benchmarking study (Zufferey *et al*., 2018), we evaluate TADs according to their enrichment of expected structural proteins, CTCF, RAD21, and SMC3, around TAD boundaries. This metric is a fold change between peaks around the boundary vs background regions. More formally, given a structural protein profile (SPP, including a number of peaks in 5 Kb intervals), we calculate ‘peak’ as the average SPP within a region surrounding boundary (*±*100 Kb) and ‘background’ as average SPP in two regions spanning 100 Kb each and located 400 Kb apart from the boundary (Zufferey *et al*., 2018). Finally, we calculate fold change 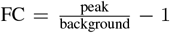 so FC greater than 0 indicates more enrichment of structural proteins around the boundary.

## 3 Results

### 3.1 Mean-variance relationships differ significantly across Hi-C experiments

As expected, we found that the variance of observations between two biological replicates varies among genomic bin pairs and depends on the magnitude of observations (Fig. 1b). We found this dependency is not similar in different experiments. For example, it depends on sequencing depths, the level of biological heterogeneity between two replicates, etc. Our data-driven variance-stabilizing approach, VSS-Hi-C, aims to consider experiment-specific mean-variance relationships to derive a transformation.

To assess the experiment-specific mean-variance relationship, we calculated mean and standard deviation (stdev) of observations as discussed in section 2.3 for Hi-C experiments from three cell lines, GM12878, IMR90, and K562, each of them including two biological replicates (Fig. 1b). We observe the expected pattern of increase of variance with increasing the average interaction frequency in all three experiments. However, the relationships differ significantly between cell lines. This implies that each experiment requires a different variance-stabilizing transformation (Fig. 1c) and thus a data-driven approach is required.

### 3.2 VSS-Hi-C signals have a unit variance

Next, we compared the variance instability (VI) of signals after different transformations for all three experiments. The lower value of this metric indicates a more stabilized variance of transformed signals. We found that VSS-Hi-C results in lower VI in all three experiments followed by another data-driven transformation, VST (Fig. 2c). These results indicate that a data-driven strategy is required for a variance-stabilization task and they outperform heuristic-based approaches with strict distributional assumptions like log, asinh and hft significantly. A VSS-Hi-C (log converge) is a linear transformation of VSS-Hi-C signals to converge to log function asymptotically and its VI does not change because this metric is invariant to the linear transformation.

To further explore the mean-variance relationship after transformations, we visualized the mean and standard deviation of signals after transformation for the best two approaches in Fig. 2a and Fig. 2b and other transformations in Fig. S1. We observe that the standard deviation of VSS-Hi-C signals is around 1 for all signal intensities as opposed to VST signals with lower standard deviation. We should note that the absolute value of variances across the whole range of signals is important in addition to VI which is a variance of variances because it implies the amount of information we keep from the original signal. To emphasize the importance of the absolute value of variances, consider a constant transformation function *t*(*x*) = *c*. This transformation has VI equal to 0 which is minimal while it ignores all the information from the original signal. Similarly, VSS-Hi-C (log converge) has a similar VI as VSS-Hi-C, however, the standard deviation of its signals is less than VSS-Hi-C (Fig. S1e). This can affect downstream analyses by putting less weight on signal differences.

One advantage of signals with a unit variance is their visualization because the changes in signal intensities have the same meaning across the range of signal intensities. A typical first step of Hi-C analysis is a heatmap visualization of interaction frequencies. It is impractical to visualize a multi-scale organization of a chromosome with a linear color scale as it only distinguishes diagonal vs. off-diagonal interactions (Fig. 1e). Therefore, a log transformation is typically applied before visualization. However, as shown previously, log signals do not have constant variance which is ideal for linear color scale. We compared the visualization of log and VSS-Hi-C signals corresponding to intra-chromosomal interactions within a 12 Mb region. Both long- and short-range interactions are better distinguished in the VSS-Hi-C heatmap. However, long-range interactions in the log heatmap are blurry as the standard deviation of low signals is higher than expected (Fig. S1a).

### 3.3 VSS-Hi-C signals improve the performance of subcompartment callers

To evaluate the utility of transformed Hi-C data for annotating subcompartments, we apply two clustering methods, the Hidden Markov model (HMM) and KMeans on raw and transformed odd-even inter-chromosomal contact matrices (section 2.5.2). To evaluate called subcompartments, we hypothesize that positions in the same subcompartment should exhibit similar activity and measure their adherance to this hypothesis with the previously-described variance explained (VE) metric (Methods). Since researchers have reported different numbers of subcompartments, we show results for *K* = *{*3, 5, 7, 9*}*. We use Hi-C data for GM12878 cell line (Rao *et al*., 2014) in this experiment because subcompartment annotation requires high-resolution Hi-C data. For data-driven transformations, VSS-Hi-C and VST, we use an inter-chromosomal contact matrix between chromosomes 1 and 2 to fit the mean-variance relationship and derive a transformation function. Then, we apply a transformation function to this and the rest of the matrices. For Haar-Fisz and ChromoR transformations, we transform data from all pairs of chromosomes together.

We observe that VSS-Hi-C (along with log and asinh transformations) outperform other transformations (Fig. 3, Fig. S2). To better understand the impact of variance stability on this task, we calculated the variance instability (VI) metric for transformed inter-chromosomal data. We observe that VI for log and asinh signals is similar to VSS-Hi-C signals (Fig. S3a) as opposed to intra-chromosomal data (Fig. 2). This might be due to the sparsity of inter-chromosomal data. Note that the robustness of estimated means, variances and their relationships in VSS-Hi-C depends on the reliability of the ordering based on another replicate. VSS-Hi-C can overestimate the variances when replicates are sparse and Aux read counts within a bin are less probable to be drawn from the same distribution. Therefore, VSS-Hi-C becomes slightly less advantageous in stabilizing the variance of inter-chromosomal data. However, it still performs comparably to log and asinh and outperforms other transformations.

**Fig. 3:**
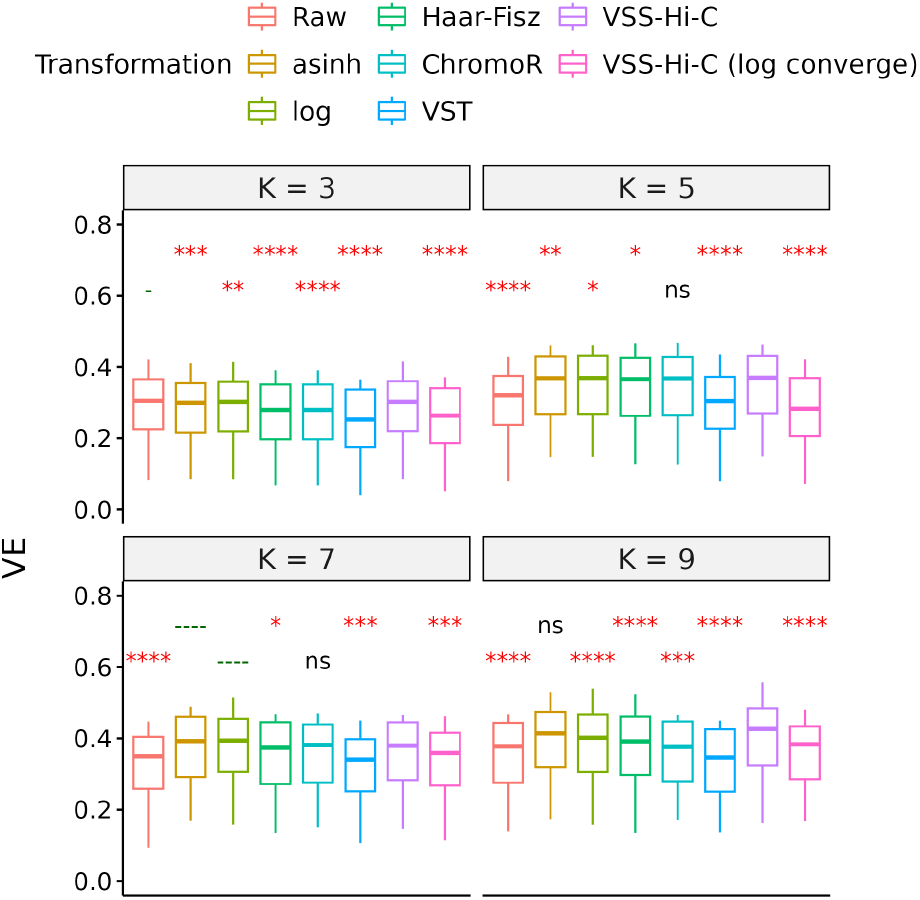
Variance explained (VE) of epigenomic features given subcompartment annotations for odd chromosomes by applying HMM on raw and transformed odd-even inter-chromosomal matrix. K indicates the number of subcompartment types in the annotation. Each box represents 12 VEs corresponding to 12 epigenomic features for a specific transformation and K. This experiment is on the GM12878 cell line. Red asterisks indicate the significance of paired one-sided t-tests with an alternative hypothesis: VSS-Hi-C signals have higher VE compared to other transformed signals. Green dashes indicate the significance of tests with a reversed alternative hypothesis: VSS-Hi-C signals have lower VE compared to other transformed signals. Non-significant (ns) labels represent (*p >* 0.05) for both tests. 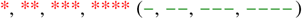 represent *p* less than 0.05, 0.01, 0.001, and 0.0001 respectively.

Furthermore, we found that Haar-Fisz and ChromoR signals have the smallest variance instability while not improving the subcompartment annotations. This is probably because of the type of this transformation that is not a monotonic function as other transformations. Therefore, two genomic bin pairs with the same interaction frequency can have different transformed signal values (Fig. S3b), and the intensity of transformed signal values becomes less informative.

### 3.4 Variance stabilization preserves the performance of TAD callers

To evaluate the utility of transformed Hi-C data for identifying topologically associating domains (TADs), we apply three TAD callers, TopDom, HiCseg, and SpectralTAD on raw and transformed data with different transformations, and calculate the fold-change metric representing the enrichment of the expected structural proteins around TAD boundaries (section 2.5.2). We found that the enrichment of structural proteins around TAD boundaries identified by SpectralTAD (Fig. 4a) and TopDom (Fig. S4a) does not change after VSS-Hi-C transformation. Further comparison of a fold-change across all transformations (Fig. 4b and Fig. S4) shows that all transformations except Haar-Fisz preserve the performance of TAD callers.

**Fig. 4:**
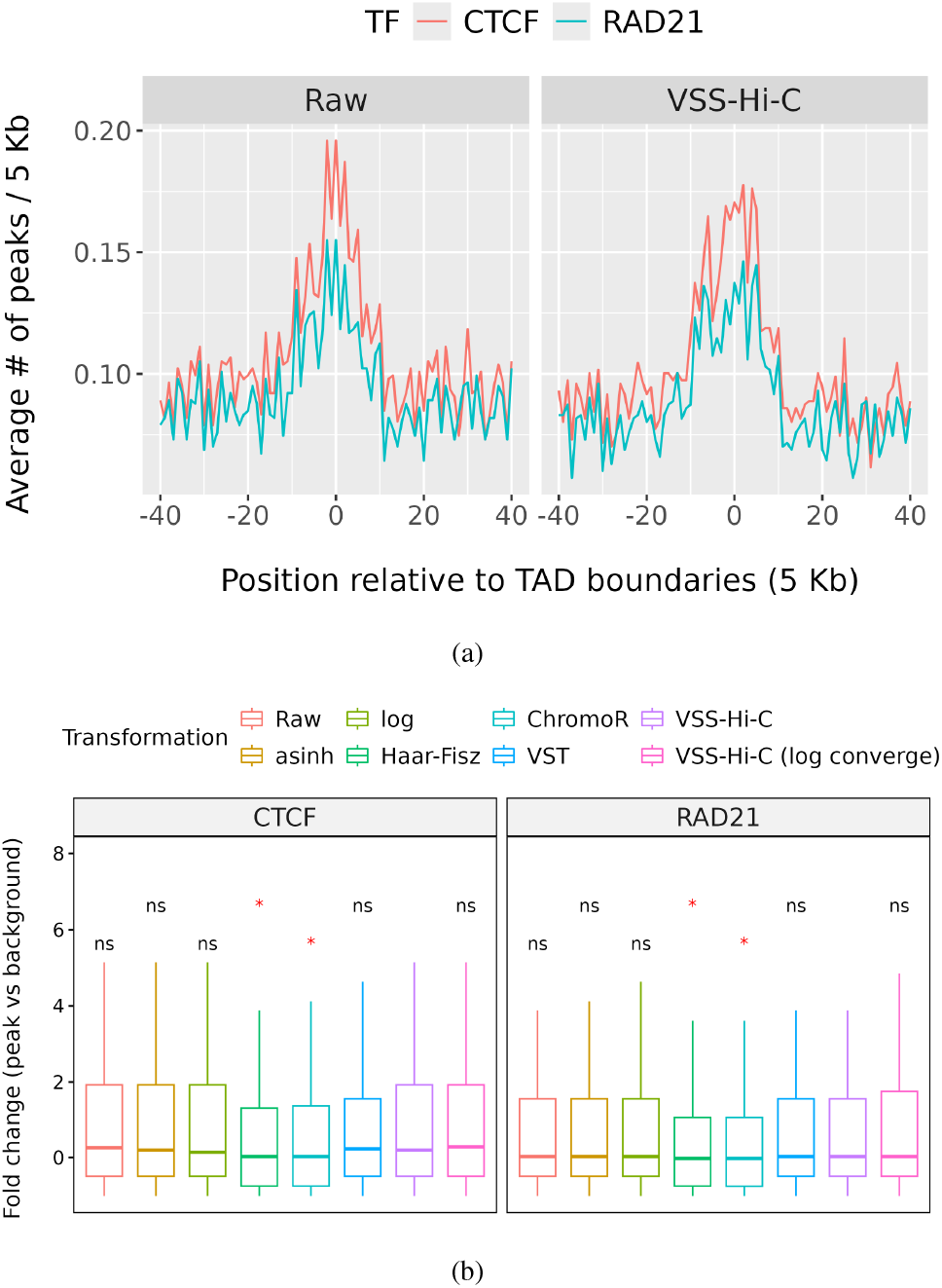
(a) Average number of ChIP-Seq peaks for CTCF and RAD21 transcription factors (TFs) per 5 Kb regions around (*±* 200 Kb) TAD boundaries identified by applying SpectralTAD on raw and VSS-Hi-C signals. (b) Fold change of CTCF and RAD21 peaks around TAD boundaries to distant regions for TADs identified by applying SpectralTAD on different transformed signals. Red asterisks indicate the significance of one-sided t-tests with an alternative hypothesis: VSS-Hi-C signals have higher fold change compared to other transformed signals. 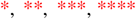 represent *p* less than 0.05, 0.01, 0.001, and 0.0001 respectively. ‘ns’ represents non-significant (*p* greater than 0.05). These plots are for IMR90 cell line.

Given a goal of variance-stabilized signals to fit the Gaussian assumption of statistical models, we expected the performance of HiCseg to increase after data transformation. However, we observed a small number of big TADs identified by HiCseg after variance-stabilizing transformations and they are not enriched with structural proteins (Fig. S5a). HiCseg is a dynamic programming (DP) algorithm that recursively calculates the likelihood of a segmentation. The analysis of a likelihood vs. the number of change points or boundaries shows that the optimum number of boundaries for variance-stabilized data is small (Fig. S5b). This is due to the assumption in HiCseg that all off-diagonal blocks (between-TAD interactions) have the same expected interaction frequency, *μ*_0_. Consequently, with transformed data, the squared error corresponding to between-TAD interactions (difference between entries in off-diagonal blocks and a constant *μ*_0_) is not negligible compared to the squared error corresponding to within-TAD interactions. Therefore, DP chooses larger TADs, less inter-TAD interactions, and less penalty from between-TAD interactions. Note that this is not a problem with raw data since the squared error for between-TAD interactions is negligible compared to the squared error for within-TAD interactions with large intensities and variances.

In general, we should pay attention to the assumptions of computational tools before inputting transformed data. Our experiments show that variance stabilization does not increase or decrease the performance of TAD callers without restrictive assumptions. However, a constant variance property can be uninteresting when the performance of a model depends on a lower variance for lower off-diagonal interaction frequencies.

## 4 Discussion and Conclusion

Here, We presented the VSS-Hi-C pipeline, a transformation approach for data-driven variance stabilization of chromatin contact counts from Hi-C data. Our results show that VSS-Hi-C signals have a unit variance across a whole range of signal intensities that are more meaningful and interpretable and provide clearer visualization of multi-scale chromatin organization. Furthermore, a constant variance property improves the performance of clustering methods relying on Gaussian observed variables for identifying subcompartments. Also, it does not degrade the performance of most computational tools developed for raw Hi-C data.

VSS-Hi-C is an extension of VSS (Bayat and Libbrecht, 2021) proposed for variance stabilization of ChIP-Seq assays to Hi-C assay. Our open-source R implementation works with a well-developed class (InteractionSet) and file format (.cool) for Hi-C data. In addition to Hi-C-specific interfaces, we included variance-stabilizing transformation functions taking two-column data frame (corresponding to two replicates of the same sample) as input to stabilize the variance of signals from other modalities.

The importance of variance stabilization has been proved for analyses including multiple samples like sample clustering (Love *et al*., 2014; Hafemeister and Satija, 2019) and differential expression analysis (Law *et al*., 2014). To our knowledge, the value of variance-stabilized signals for analysis tasks within a sample is not explored widely. Here, We showed two Hi-C analysis tasks benefiting from signals with constant variance, visualization, and calling subcompartments. Exploring other tasks related to Hi-C data or other modalities that benefit from variance-stabilized signals is an interesting future work. For example, loop callers that rely on the enrichment of interaction frequencies compared to neighbour genomic bin pairs might benefit from VSS-Hi-C signals to call significant loops.

Variance instability is one factor resulting in less interpretable and meaningful Hi-C signals. There are other bias factors hiding true signal strengths, for example, genomic biases resulting in non-uniform visibility of genomic bins. Different normalization approaches have been proposed to solve inequal visibility of genomic bins in Hi-C data (Imakaev *et al*., 2012; Hu *et al*., 2012). A method to both normalize and stabilize the variance of read counts is another interesting future work.

