## Supplementary Information for "VSS-Hi-C: Variance-stabilized signals for chromatin contacts"

### VSS-Hi-C: Variance-stabilized signals for chromatin contacts (supplementary information)

Table. S 1: Data sources.

| Data type | Cell type | link |
|---|---|---|
| ChIP-Seq data | GM12878 | CTCF: ENCFF710VEH, ENCFF473RXY, ENCFF833FTF, ENCFF002DAJ<br>RAD21: ENCFF002CPK, ENCFF753RGL<br>SMC3: ENCFF686FLD<br>HMs and DNase-Seq: 'E116-*.bigwig' files |
| | K562 | CTCF: ENCFF002CEL<br>RAD21: ENCFF002CXU<br>SMC3: ENCFF041YQC |
| | IMR90 | CTCF: ENCFF453XKM<br>RAD21: ENCFF195CYT |
| Hi-C data | GM12878 | Biological replicate 1: HIC001-HIC018<br>Biological replicate 2: HIC019-HIC029 |
| | K562 | Biological replicate 1: HIC069-HIC070<br>Biological replicate 2: HIC071-HIC074 |
| | IMR90 | Biological replicate 1: HIC050-HIC054<br>Biological replicate 2: HIC055-HIC056 |

### 1 Alternative variance-stabilizing transformations

#### 1.1 Haar-Fisz transformation

Haar-Fisz transformation Gaussianizes and stabilizes the variance of the inhomogeneous one-dimensional Poisson process [Fryzlewicz and Nason(2004)Fryzlewicz and Nason] using a modified version of forward and inverse Haar discrete wavelet transform (DWT).

The forward Haar DWT works as follows: given an input vector $v = (v_0, v_1, ..., v_{N-1})$ of size $N$, where $N$ is a power of 2 ($N = 2^J$), they define $s^0 = v$ and recursively perform the following steps:

$$s^j = (s_0^j, s_1^j, ..., s_{2^{J-j}-1}^j),\ s_i^j = \frac{s_{2i}^{j-1} + s_{2i+1}^{j-1}}{2}, \tag{1}$$

$$d^j = (d_0^j, d_1^j, ..., d_{2^{J-j}-1}^j),\ d_i^j = \frac{s_{2i}^{j-1} - s_{2i+1}^{j-1}}{2} \tag{2}$$

for $j = 1, ..., J$. The elements of $s^j$ and $d^j$ represent the smooth and detail of the original vector $v$ at scale $2^j$. The detail vectors from all scales and the smooth element from the coarsest scale, $(s^J, d^J, d^{J-1}, ..., d^1)$, are enough to reconstruct the original vector $v$.

The inverse Haar DWT reverses the equations 1 and 2 to give:

$$s_{2i}^{j-1} = s_i^j + d_i^j, s_{2i+1}^{j-1} = s_i^j - d_i^j, \tag{3}$$

for $i = 0, ..., 2^{J-j} - 1$ where $j = J, ..., 1$. And, reconstructed smooth elements at the first layer are the original vector ($v = s_0$).

The Haar-Fisz transformation is very similar to the Haar DWT with the modification that at each scale $j$, it defines $f^j$ after calculating $s^j$ and $d^j$ by

$$f_i^j = \begin{cases} 0 & \text{if } s_i^j = 0, \\ d_i^j/\sqrt{s_i^j} & \text{otherwise.} \end{cases} \tag{4}$$

And, inverse Haar DWT is applied on $(s^J, f^J, f^{J-1}, ..., f^1)$ rather than $(s^J, d^J, d^{J-1}, ..., d^1)$.

The reason that the Haar-Fisz transformation can stabilize the variance is based on the Fisz theorem [Fisz(1955)Fisz] that guarantees the variance of 1 for a specific function of two Poisson random variables with large and close expectations. The new reconstruction function of smooth elements using $f$ elements is equivalent to that specific function. Therefore, if neighborhood elements in the smooth vectors are close to each other, the Haar-Fisz transformation guarantees a stabilized variance for the reconstructed vector.

We should sort the inhomogeneous chromatin contact counts to force the neighbor elements to be close to each other. Next, we should pad it with 0 to the length of a power of 2. We can apply the Haar-Fisz transformation to the processed vector to stabilize its variance. We used 'hft' function from 'haarfisz' package in R to apply this transformation.

The disadvantage of Haar-Fisz transformation is its Poisson assumption about the distribution of the input vector. Therefore, data-driven Haar-Fisz transformation was proposed where the calculation of $f$ in equation **??** is replaced with

$$f_i^j = \frac{d_i^j}{2h^{1/2}(s_i^j)}, \tag{5}$$

where $h$ is estimated from data.

#### 1.2 Variance-stabilizing transformation using estimated dispersions from DESeq2 (VST)

The DESeq2 [Love *et al.*(2014)Love, Huber, and Anders] method models within-group variability of gene $i$ by the dispersion parameter $\alpha_i$, which describes the variance of counts via $\text{Var}(K_{ij}) = \mu_{ij} + \alpha_i \mu_{ij}^2$, where $K_{ij}$ and $\mu_{ij}$ are observed and expected read counts for gene $i$ in sample $j$. Accurate estimation of $\alpha_i$

is not possible when number of samples is not large. Therefore, they employ empirical Bayes shrinkage for dispersion estimation. Their procedure has 3 steps. First, they estimate gene-wise dispersions using maximum likelihood given observations for each gene. Then, they assume that there is a relationship between gene expression abundance and dispersion such that genes with similar average expression have similar dispersion. Therefore, they learn a trend between average gene expression and estimated dispersions from the first step. This provides an accurate estimate of the expected dispersion given the average expression of a gene. To account for both gene-wise dispersion estimations and expected one given a trend, they calculate final dispersions by shrinking gene-wise estimation towards the trend. The final dispersions together with mean expressions provide estimations of the variances ($\mathrm{Var}(K_{ij}) = \mu_{ij} + \alpha_i \mu_{ij}^2$). Then, variance-stabilizing transformation is obtained by numerical integration (equation 4 in the main manuscript) of the spline fitted to the estimated mean and variance pairs.

#### 2 Supplementary figures

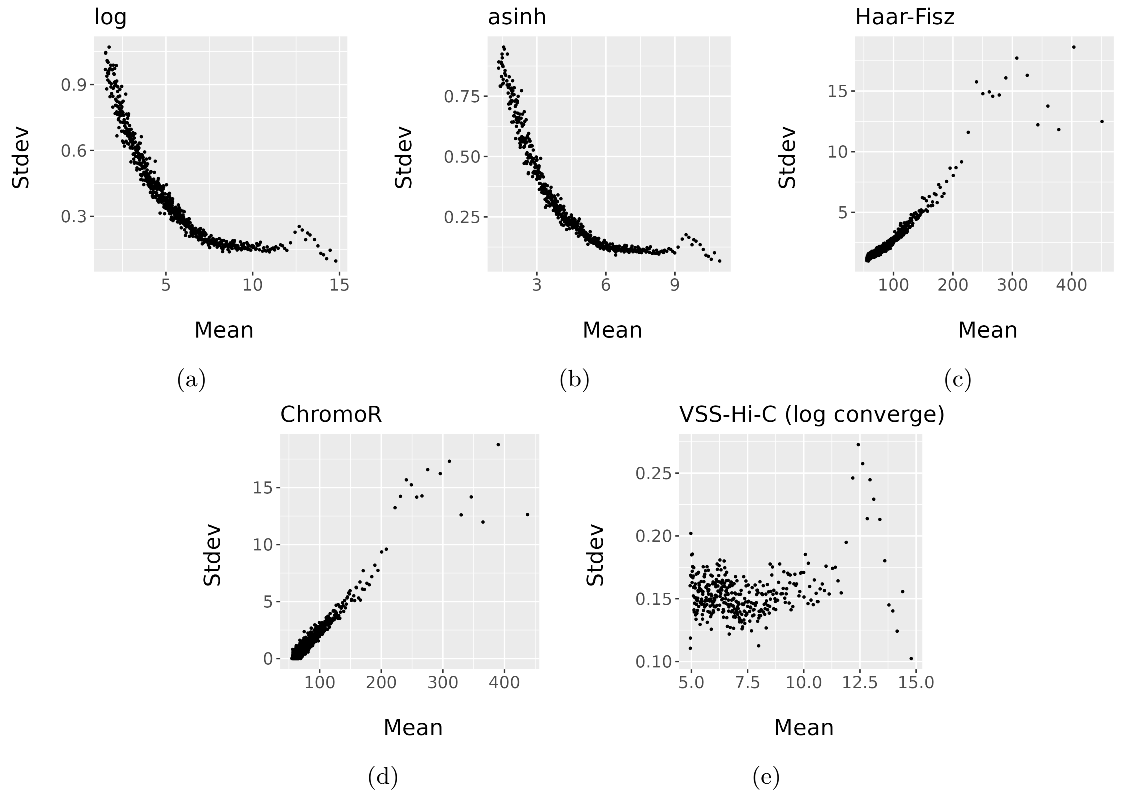


Fig. S1: Standard deviation (stdev) vs mean for transformed signals. These plots are for chromosome 13 of the GM12878 cell line.

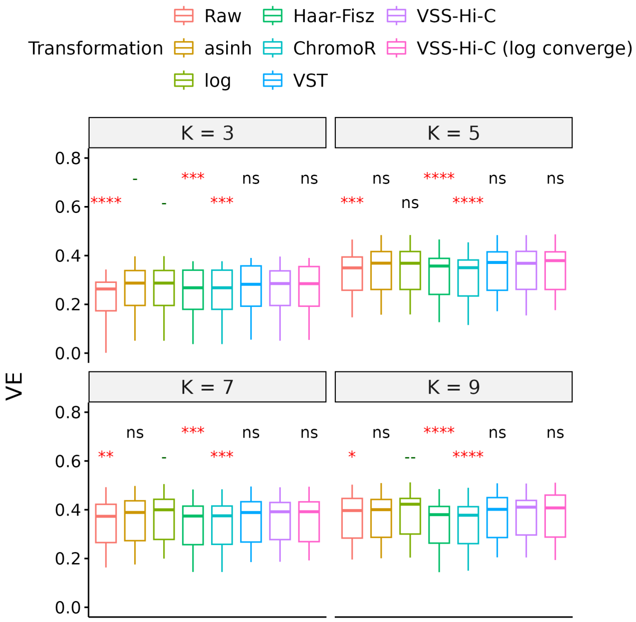


(a)

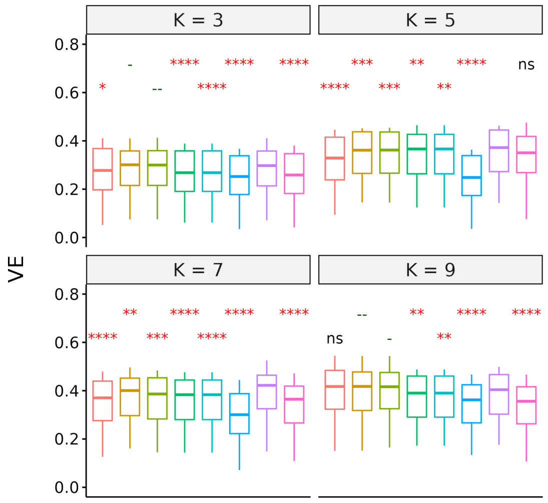


(b)

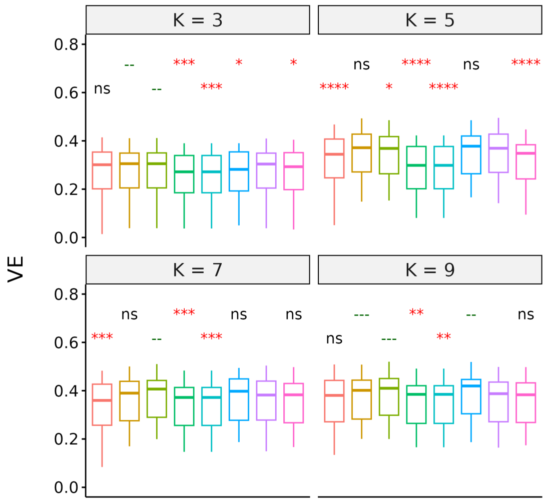


(c)

Fig. S2: Variance explained (VE) of epigenomic features given subcompartment annotations for (a) even chromosomes by applying HMM, (b) odd chromosomes by applying KMeans, and (c) even chromosomes by applying KMeans, on raw and transformed odd-even inter-chromosomal matrix. K indicates the number of subcompartment types in the annotation. Each box represents 12 VEs corresponding to 12 epigenomic features for a specific transformation and K. This experiment is on the GM12878 cell line. Red asterisks indicate the significance of paired one-sided t-tests with an alternative hypothesis: VSS-Hi-C signals have higher VE compared to other transformed signals. Green dashes indicate the significance of tests with a reversed alternative hypothesis: VSS-Hi-C signals have lower VE compared to other transformed signals. Non-significant (ns) labels represent ($p > 0.05$) for both tests. *, **, ***, **** (-, --, ---, ---- ) represent $p$ less than 0.05, 0.01, 0.001, and 0.0001 respectively.

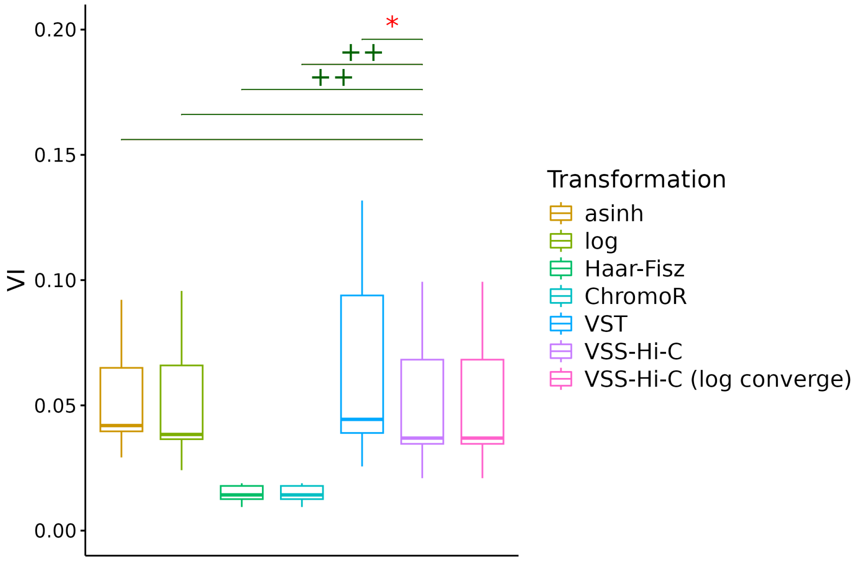


(a)

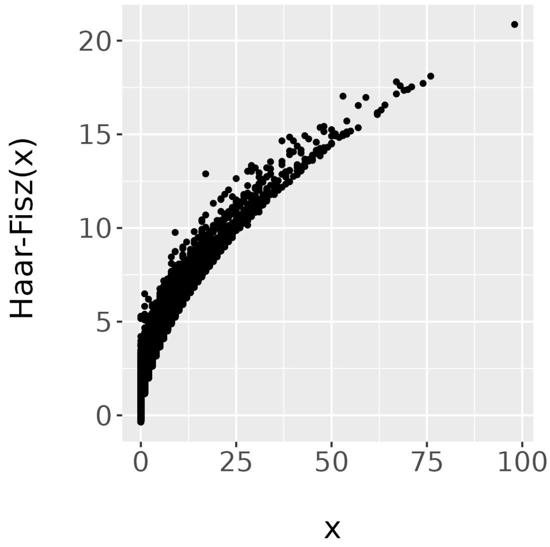


(b)

Fig. S3: (a) Variance instability (VI) of transformed inter-chromosomal Hi-C data. Red asterisks indicate the significance of paired one-sided t-tests with an alternative hypothesis: VSS-Hi-C signals have less VI compared to other transformed signals. Green pluses indicate the significance of paired one-sided t-tests with an alternative hypothesis: VSS-Hi-C signals have greater VI compared to other transformed signals. Non-significant labels ($p > 0.05$) are removed. *, **, ***, **** (+, ++, +++, ++++) represent $p$ less than 0.05, 0.01, 0.001, and 0.0001 respectively. (b) Haar-Fisz vs raw signals for inter-chromosomal Hi-C data.

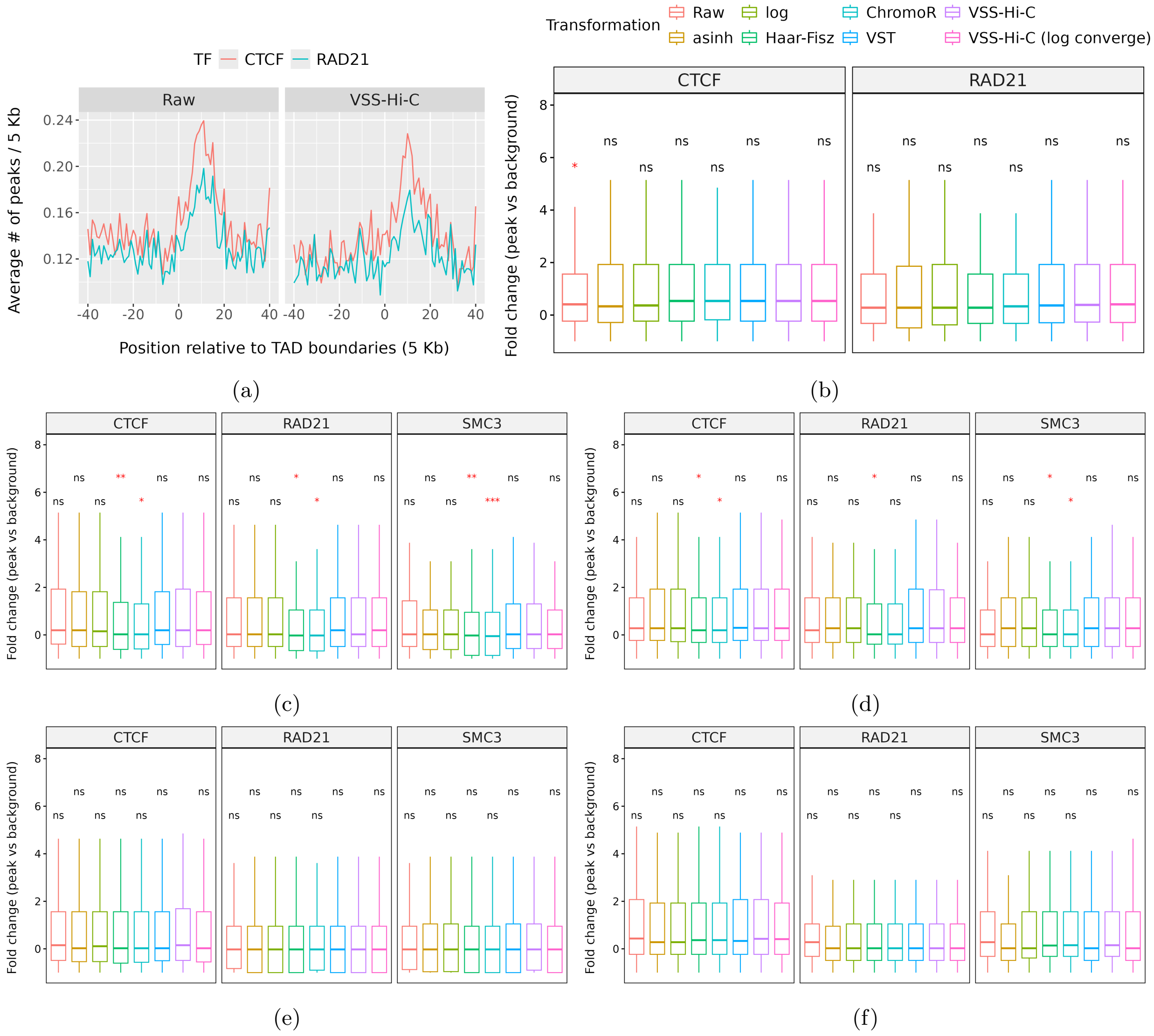


Fig. S4: (a) Average number of ChIP-Seq peaks for CTCF and RAD21 transcription factors (TFs) per 5 Kb regions around (± 200 Kb) TAD boundaries identified by applying TopDom on raw and VSS-Hi-C signals (IMR90 cell line). (b-f) Fold change of CTCF, RAD21, and SMC3 peaks around TAD boundaries to distant regions for TADs identified by applying TopDom on (b) IMR90, (d) GM12878, (f) K562 and applying SpectralTAD on (c) GM12878 and (e) K562 cell lines. Red asterisks indicate the significance of one-sided t-tests with an alternative hypothesis: VSS-Hi-C signals have higher fold change compared to other transformed signals. *, **, ***, **** represent $p$ less than 0.05, 0.01, 0.001, and 0.0001 respectively. 'ns' represents non-significant ($p$ greater than 0.05).

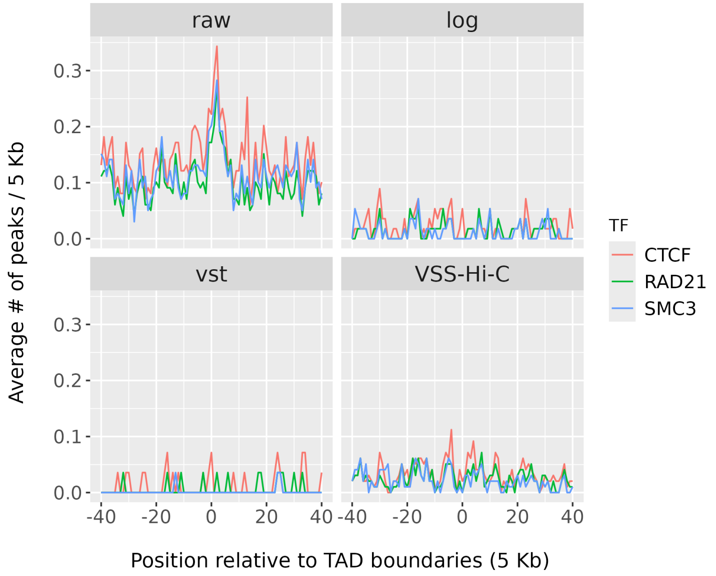


(a)

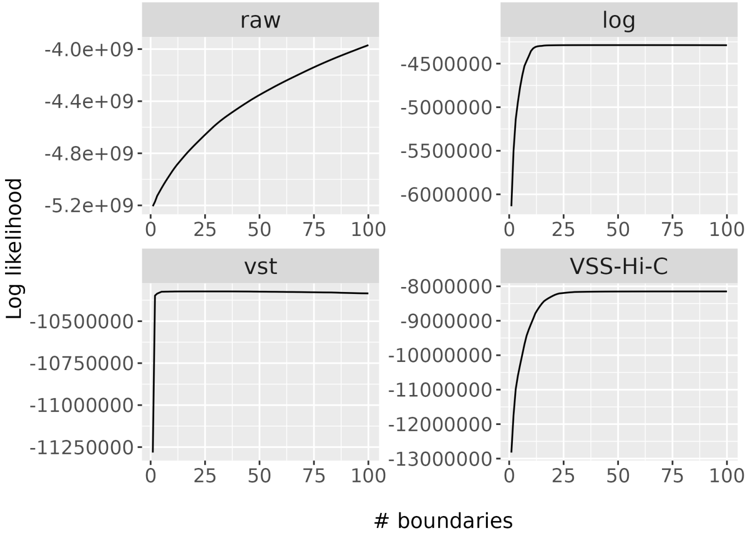


(b)

Fig. S5: (a) Average number of ChIP-Seq peaks for CTCF, RAD21, and SMC3 transcription factors (TFs) per 5 Kb regions around (± 200 Kb) TAD boundaries identified by applying HiCSeg on raw and transformed signals. (b) Log-likelihood of HiCSeg model vs number of change points or boundaries for raw and transformed signals. These plots are for GM12878 cell line.
